# Stability of dynamic functional architecture differs between brain networks and states

**DOI:** 10.1101/533307

**Authors:** Le Li, Bin Lu, Chao-Gan Yan

**Affiliations:** CAS Key Laboratory of Behavioral Science, Institute of Psychology, Beijing, China; Department of Psychology, University of Chinese Academy of Sciences, Beijing, China; Magnetic Resonance Imaging Research Center, Institute of Psychology, Chinese Academy of Sciences, Beijing, China; Department of Child and Adolescent Psychiatry, NYU School of Medicine, New York, NY, USA

## Abstract

Stable representation of information in distributed neural connectivity is critical to function effectively in the world. Despite the dynamic nature of the brain’s functional architecture, characterizing its temporal stability has been largely neglected. Here we characterized stability of functional architecture for each brain voxel by measuring the concordance of dynamic functional connectivity (DFC) over time, and explored how stability was modified by movie watching. High-order association regions, especially the default mode network, demonstrated high stability during resting state scans, while primary sensory-motor cortices revealed relatively lower stability. During movie watching, stability in the primary visual cortex was decreased, which was associated with larger DFC variation with neighboring regions. By contrast, higher-order regions in the ventral and dorsal visual stream demonstrated increased stability. The distribution of functional stability and its modification describes a profile of the brain’s stability property, which may be useful reference for examining distinct mental states and disorders.

---

Stability is a critical feature for consciousness, to maintain stable and consistent representation of information by distributed neural activity and connectivity patterns over time ^1^. The brain coordinates information from multiple regions and moments through distributed functional connections among regions in conscious states ^2, 3^, thus a stable functional architecture is essential. However, despite the neurobiological significance of such stability, how stability is distributed across brain systems and how it is modified when executing tasks remain largely unknown.

The brain implements cognitive functions in a spatially organized way ^2, 4^. The association regions, involved in high-order cognitive processing, are more globally connected, compared to unimodal regions that underlie primary sensory-motor processing, from a static perspective ^5, 6^. From a dynamic perspective, studies report higher temporal variability in association areas in terms of functional connectivity with other regions, while lower temporal variability is found in unimodal areas in the resting state ^7, 8^. This is consistent with the hypothesis that association regions switch or change their functional connections frequently since they integrate information from various modalities into multimodal representations ^9^, thus exhibiting a lower level of stability of functional architecture. However, competing evidence and hypotheses exist. Between-session intra-subject functional connectivity variability was shown to be smaller in association regions than unimodal regions ^10^. In addition, association regions were proposed to process information over a longer time scale (in minutes) than unimodal regions (in seconds) ^11^. Therefore, association regions may serve as hubs to coordinate neural signals over time, and would be hypothesized to display high stability of functional architecture which requires direct confirmation. Studies examining flexibility ^7, 8^ could have failed to support the alternate hypothesis due to two factors: 1) they characterized functional architecture with the Automated Anatomical Labeling (AAL) atlas, a structural atlas that is considered coarse and functionally inaccurate, and cannot adequately reflect the functional architecture of the human brain ^12^; and 2) by omitting quantification of stability as a property, emphasizing flexibility may highlight areas with low signal-to-noise ratio, e.g., anterior temporal regions. Thus, it is crucial to test the two competing hypotheses empirically to enhance our understanding of the dynamic architecture of human brain, by precisely characterizing the stability of functional architecture voxel-by-voxel.

To implement a specific task, the brain’s functional architecture changes according to the current task demands of cognitive processes ^13, 14, 15^. This change in turn results in modification in the stability of functional architecture. Cole et al. (2013) showed high between-task flexibility of functional architecture for the frontoparietal network ^13^, while the stability within a continuous task (e.g., a naturalistic task) remains unknown. Movie watching, for example, requires viewers to constantly integrate presented stimuli which are closely related to each other in context over time. Prior studies with naturalistic tasks have revealed dynamic changes of functional connectivity of the default mode network (DMN) that was specifically induced by the task ^16^. However, the stability profile in such a real-life situation remains unknown. Integration of visual and auditory information involves the occipital temporal cortex (OTC) and superior temporal sulcus (STS) ^17, 18^, which can be regarded as association regions for this task. Functional stability of these regions should be increased due to the need to constantly integrate information over a long time scale in natural viewing tasks, though this hypothesis needs to be tested.

Here we sought to precisely characterize stability of functional architecture across the brain and its modification during task states. Resting-state fMRI can measure the “intrinsic” brain functional architecture which is consistently present across a wide variety of cognitive states ^4, 19^. We first analyzed resting-state data to quantify stability of functional architecture in its intrinsic form across the brain. We defined stability of functional architecture for a brain voxel as the concordance of its voxel-level dynamic functional connectivity (DFC) over time. Furthermore, we explored how the stability profile was modified by a naturalistic task from its intrinsic form, through comparison of functional stability between a movie-watching task and resting state, using a movie-watching dataset.

## Results

### Profile of stability of intrinsic functional architecture

We analyzed resting-state fMRI data of 216 young adults from the CoRR (Consortium for Reliability and Reproducibility) release ^20^, to examine intrinsic functional stability across the brain. The data contained two scanning sessions acquired on different days.

Functional stability for a brain voxel was defined as the Kendall’s coefficient of concordance (KCC, also known as Kendall’s W) of DFC over time between that voxel and all other regions in the brain (Methods). DFC was calculated over consecutive segments of data in a sliding window approach ^21^. Notably, analyses were conducted in a voxel-to-voxel approach, in which the KCC of a voxel was computed based on the features of its voxel-level DFC maps (Fig. 1). Such approach can provide a refined and global characterization of how a brain region changes its functional architecture over time. The derived KCC for each subject was z-standardized across a grey matter mask. Standardization minimizes the effect of overall discrepancy in KCC across subjects and conditions, and thus enabled us to examine relative differences among brain regions ^22^. A higher KCC value for a region means its functional architecture configuration is more consistent and stable over time.

**Figure 1.**
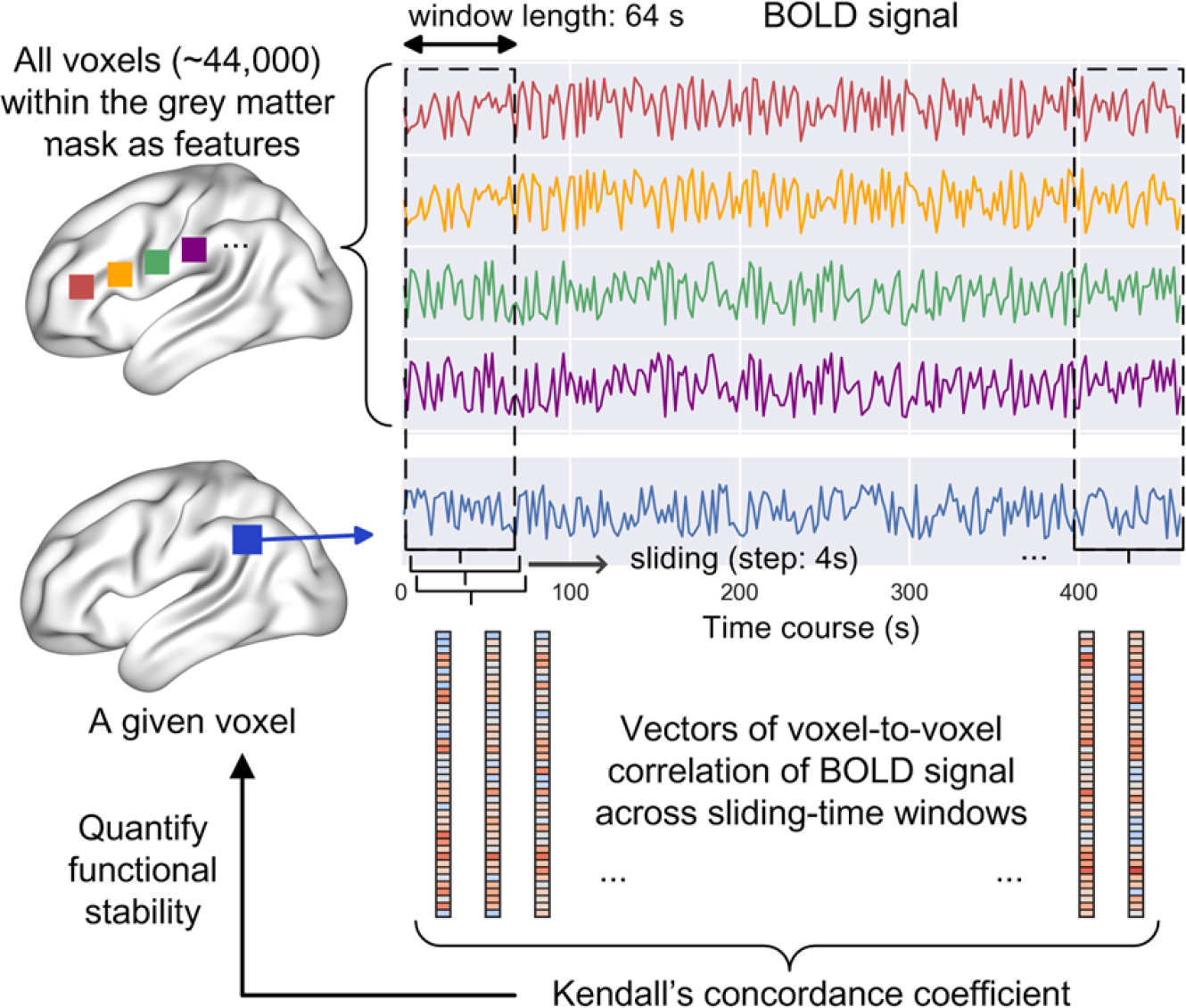
Schematic diagram shows how the stability of functional architecture is computed in a voxel-to-voxel approach. Dynamic functional connectivity (DFC) for a given voxel is calculated with all voxels within the grey matter mask for each window, and constitutes features for the functional architecture for that voxel. The rectangular windows are 64 s in length, with 4 s sliding steps. Kendall’s concordance coefficient is computed based on DFC across windows, and quantifies the functional stability for that voxel.

One-sample T-tests (n = 216) revealed that in both sessions, the intrinsic stability of functional architecture differed substantially across the brain. First, the apex of intrinsic stability was observed bilaterally in the dorsolateral prefrontal cortex (DLPFC), anterior insula (AIns), lateral temporal cortex (LTC), supramarginal gyrus (SMG), angular gyrus (AG), medial prefrontal cortex (mPFC), posterior cingulate cortex (PCC), and occipitoparietal cortex (Fig. 2A,B in red). These regions are high-order association areas. At the other extreme, the lowest intrinsic stability was found in regions near cavities and ventricles, including the anterior temporal lobe, orbitofrontal cortex, and caudate nucleus (Fig. 2A,B in blue). High susceptibility to artifacts results in low signal-to-noise ratio in these regions ^23^, which inevitably leads to substantial decrease in functional stability. Other regions showed intermediate levels of intrinsic stability. Compared to the high-order association regions, unimodal regions (including auditory, somatosensory, visual, and motor regions) displayed relatively lower intrinsic stability (Fig. 2A,B), indicating that their functional architectures were less consistent over time. Within the framework of brain networks defined by Yeo et al. ^24^, the ratio of voxels with higher stability was largest for the DMN, followed by the frontoparietal network (FPN) and the ventral attentional network (VAN) (Fig. 2C,D). Notably, the pattern of intrinsic stability across the brain was similar between the two resting-state sessions, indicating high reliability of these results. The averaged stability across all subjects resembled the T-test result (Fig. S1).

**Figure 2.**
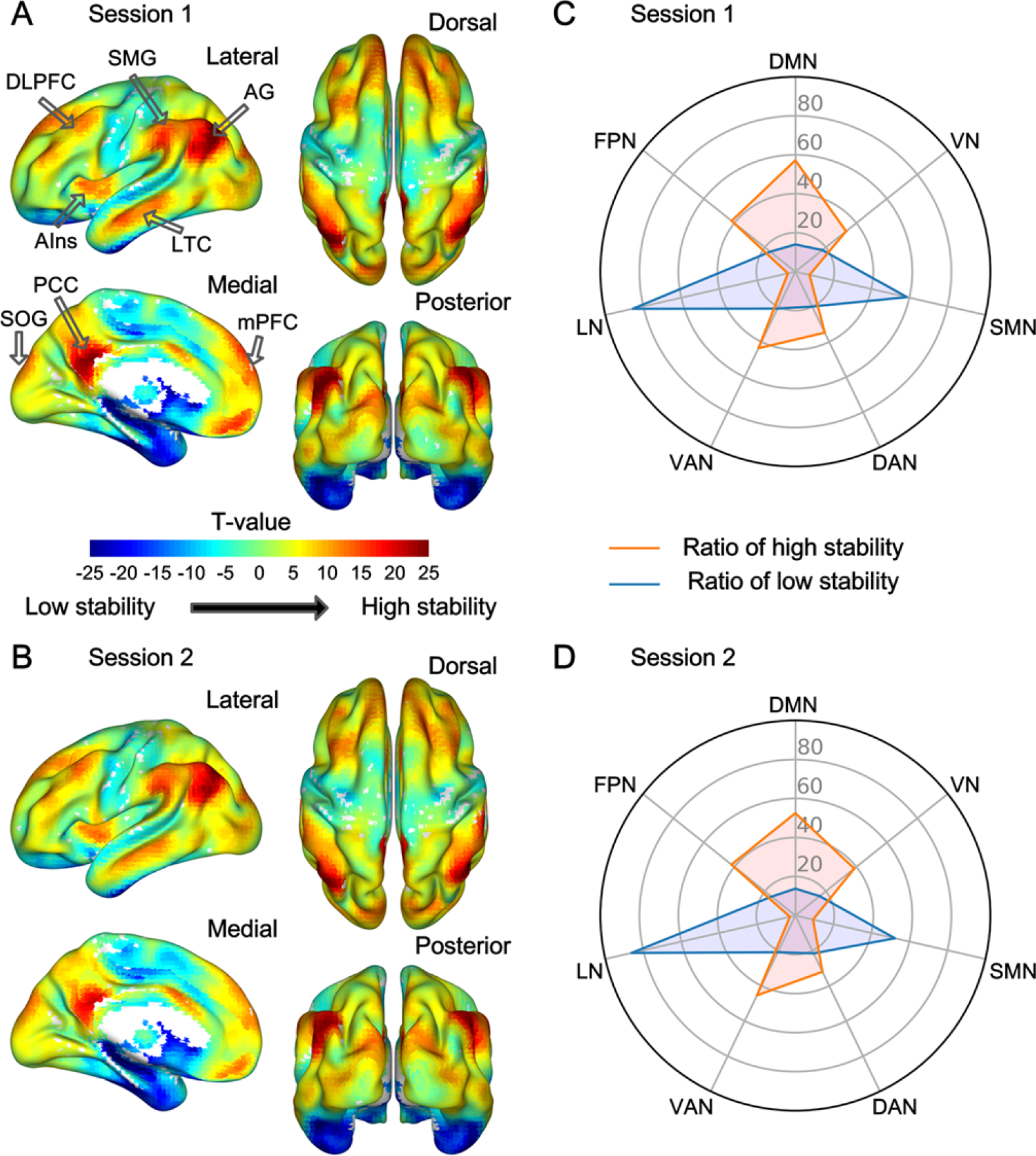
Profile of intrinsic functional stability across the brain. (A,B) Results of one-sample T-tests on functional stability (converted to z-scores) in resting state. (C,D) the ratio of voxels showing high and low stability for the seven brain networks. A positive value (in yellow to red) denotes high stability while a negative value (in cyan to blue) denotes low stability. The ratio was computed as the number of significant voxels after Gaussian random field correction divided by the total number of voxels in a network. High stability is observed in several association regions indicated by black hollow arrows. DLPFC, dorsolateral prefrontal cortex; AG, angular gyrus; AIns, anterior insular; LTG, lateral temporal cortex; SOG, superior occipital gyrus; PCC, posterior cingulate cortex; mPFC, medial prefrontal cortex.

Notably, as shown in Fig. 2, some regions in the visual network exhibited an above average level of functional stability (in yellow-orange). This observation might seem to contradict the finding that the brain’s functional architecture was more stable for association regions than for unimodal regions. We thus compared functional stability between associative and primary visual cortices (Methods). Four associative and six primary visual regions were selected, for each hemisphere (Fig. 3, see Yeo, et al. 2011 for the coordinates). Functional stability was averaged for each of the two types of visual regions, respectively, and then compared between them with paired-sample tests for each hemisphere. The results revealed that high-order association regions also exhibited higher functional stability than unimodal regions in the visual network of both the left hemisphere (t = 4.28, p < 0.001 for the first session; t = 4.65, p < 0.001 for the second session; Fig. 3) and the right hemisphere (t = 3.54, p < 0.001 for the first session; t = 4.98, p < 0.001 for the second session; Fig. 3).

**Figure 3.**
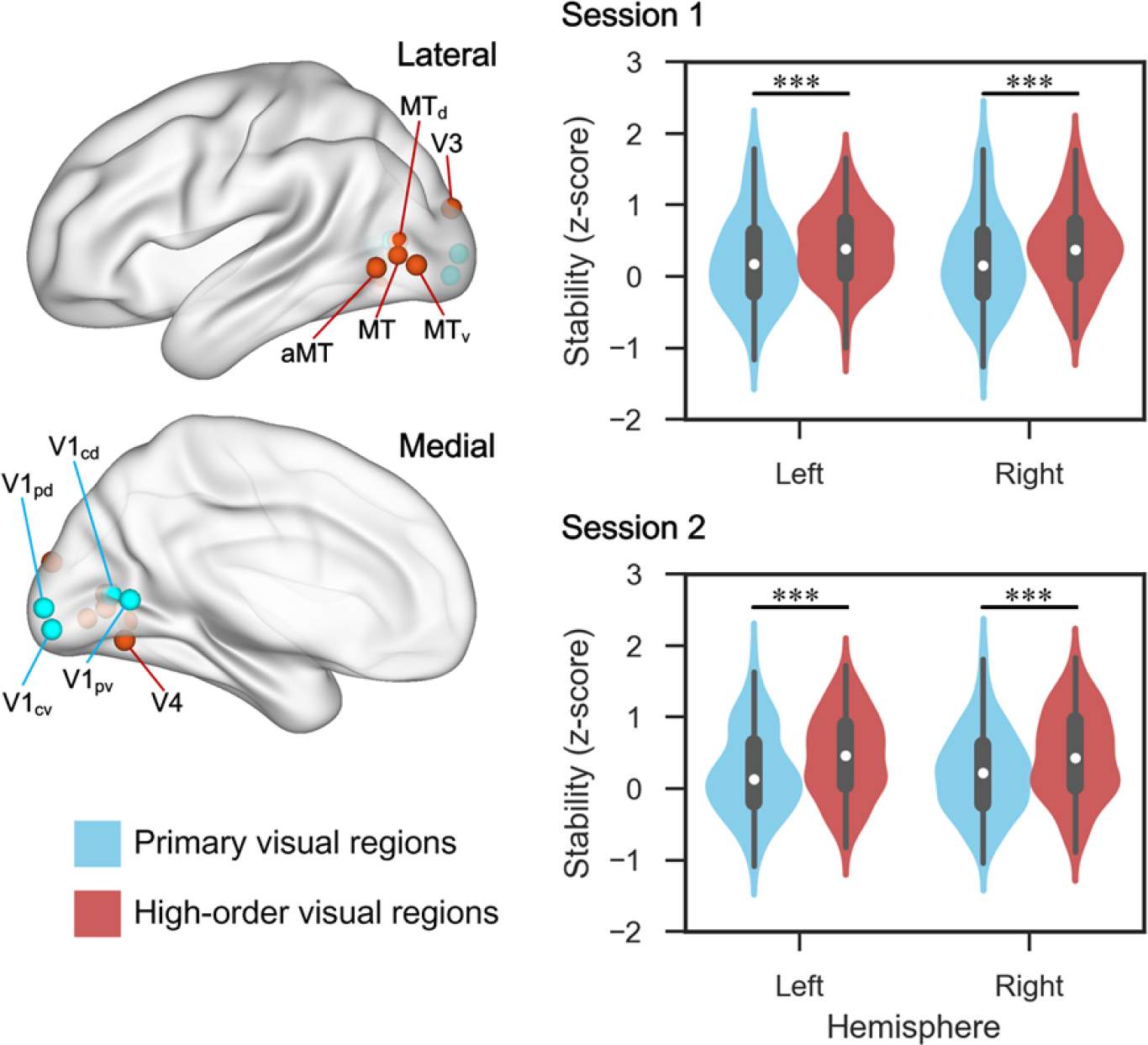
Difference of intrinsic functional stability between high-order associative visual regions and primary visual regions. The locations of presented regions of interest of these two types of regions are shown in the left panel. The violin plots in the right panel reveal the distribution and difference of functional stability between them for both hemispheres and both sessions. ***, p < 0.001. MT, middle temporal area; V1, primary visual area; V1_pd_, dorsal part of peripheral V1; p, peripheral; c, central; d, dorsal; v, ventral; a, anterior.

Furthermore, we examined whether stability exceeded random levels. Simulated data were created by randomizing the phases while maintaining the amplitudes of resting-state signals. This removed the temporal alignment of neural signals which is essential to measure stability, and thus resulted in a baseline level. Functional stability raw values were compared between the observed and simulated data with paired-sample T-tests. Results revealed that in almost all voxels across the brain, the observed functional stability was greater than the simulated functional stability (all p < E-10; Fig. S2). Taken together with the prior results, this indicates that functional stability does not exist in simulated random data, and that it is distributed across the brain in a biological meaningful way.

### Stability of functional architecture during natural viewing

We next moved to investigate functional stability of the brain in a complex naturalistic task with a continuous state. Here a movie watching task was employed, during which viewers constantly received and integrated changing audiovisual stimuli over time, to comprehend the movie. The dataset from the HBN (Healthy Brain Network) released by the Child Mind Institute ^25^ was analyzed. For this dataset, fMRI data from 32 children and adolescents were entered into analyses, consisting of two runs of 5-min resting-state scans, followed by another run of movie watching. The movie was a 10-min clip of an animated film named “Despicable Me”. We divided the movie-watching run into two halves, and then averaged functional stability between the two halves and between the two resting-state runs (Methods). The averaged functional stability was contrasted between movie watching and resting state with paired-sample T-tests. This comparison allowed us to examine how stability was modified from its intrinsic form (i.e., resting state) to a natural viewing task.

Results showed that functional stability was increased during movie watching in the bilateral occipitotemporal cortex (OTC), left posterior middle temporal gyrus (pMTG), left posterior fusiform gyrus (pFG), right posterior inferior temporal gyrus (pITG), right superior temporal sulcus (STS), and left intraparietal sulcus (IPS) (voxel-level p < 0.001, Gaussian Random Field corrected to p < 0.01, two-tailed, the same below; Fig. 4A and table 1). Most of these loci are in the higher visual processing stream. Decreased stability was observed for movie watching in the mPFC, and the expanse of bilateral medial and posterior occipital region, including the calcarine sulcus (CalS), cuneus, and lingual gyrus (LG) (Fig. 4A and table 1). Notably, the within-subject design of the contrast between movie watching and resting state can yield large effect sizes despite a small sample size ^26^.

**Table 1.** Stability difference of movie watching vs. resting state

| Cluster size | area | x | y | z | T-value |
| --- | --- | --- | --- | --- | --- |
| 285 | L.pMTG | -48 | -54 | 9 | 6.959 |
|  | L.pFG | -42 | -45 | -18 | 5.390 |
| 112 | R.OTC | 45 | -60 | -3 | 6.753 |
| 117 | L.IPS | -39 | -33 | 57 | 6.184 |
| 66 | R.STS | 42 | -30 | -3 | 6.060 |
| 53 | R.pITG | 45 | -51 | -21 | 5.028 |
| 1303 | L.calcarine | -6 | -90 | -3 | -9.449 |
|  | L.cuneus | -12 | -93 | 27 | -7.303 |
|  | R.cuneus | 18 | -90 | 24 | -7.218 |
|  | L.LG | -15 | -66 | -6 | -6.749 |
|  | R.LG | 12 | -75 | -9 | -6.341 |
|  | R.calcarine | 15 | -63 | 12 | -3.853 |
| 46 | mPFC | 9 | 57 | 9 | -5.675 |

**Figure 4.**
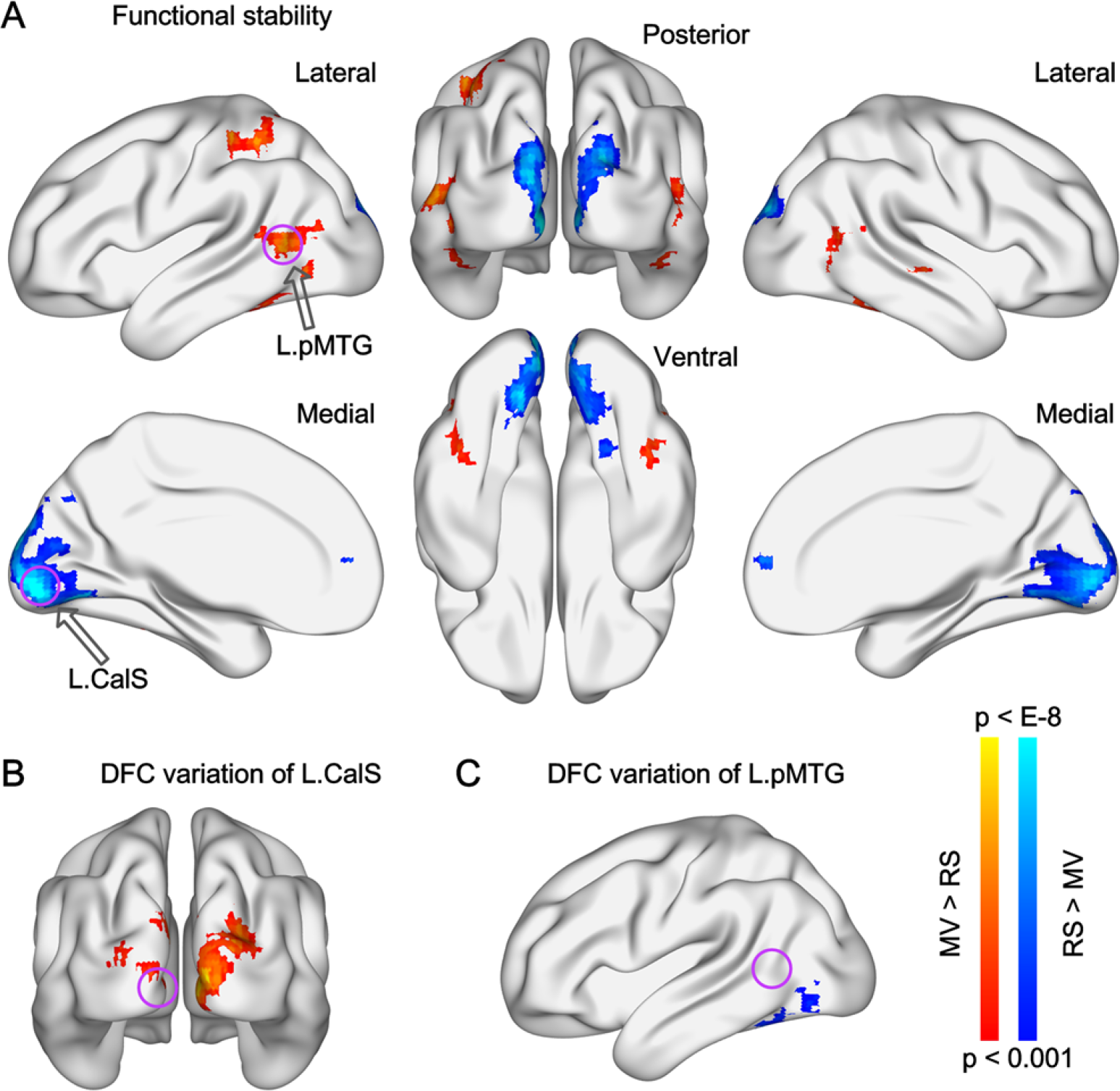
Differences of functional stability and of ROI-based DFC variation between movie watching and resting state. Brain maps of T-values show the results of paired-sample T-tests between movie watching and resting state on functional stability (A), and on DFC variation of left calcarine sulcus (L.CalS) (B) and of left posterior middle temporal gyrus (L.pMTG) (C). The location of the two seed regions are indicated by purple circles. L, left; R, right; MV, movie watching; RS, resting state.

To examine whether regions in which stability changed were actually engaged by movie watching, we conducted an analysis of inter-subject correlation (ISC) of neural activity ^27^. The ISC measures the synchronization of responses to naturalistic stimuli across subjects, which should only be caused by common cognitive processes ^28^. It can reveal which brain regions were engaged when subjects watched the movie, and is not sensitive to within-subject confounding factors. The results revealed significant ISC (r > 0.25 in average and p < 0.001 in one-sample T-test versus 0, Fig. 5) bilaterally in the occipital lobe, OTC, superior temporal cortex, occipitoparietal cortex, IPS, SMG, and precentral gyrus. These areas included all the regions in which stability was modified by movie watching, except the mPFC, suggesting that stability modification was relevant to regional engagement rather than within-subject confounding factors.

**Figure 5.**
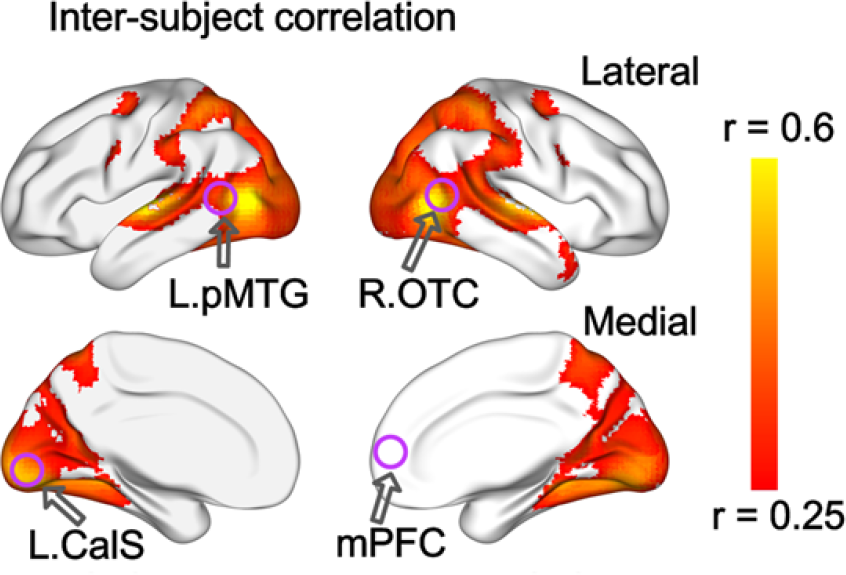
Averaged inter-subject correlation of neural activity during movie watching. The colored area was masked by a threshold of r > 0.25 and by Gaussian random field correction for multiple comparisons (p < 0.001 at voxel level and cluster p < 0.01) in one-sample T-test. The purple circles indicated four representative regions in which functional stability was modified by movie watching. L, left; R, right; pMTG, posterior middle temporal gyrus; OTC, occipitotemporal cortex; CalS, calcarine sulcus; mPFC, medial prefrontal cortex.

Functional stability of specific regions was measured based on the whole-brain DFC for those regions. A further step is to probe which connections specifically contributed to the difference in stability observed between states. To partly address this problem, we took the regions of which the stability was modified by movie watching as regions of interest (ROIs, including the left pMTG and left CalS), and then compared their DFC variation maps between movie watching and resting state. These two ROIs were selected because they were representative visual regions showing the most significant stability difference in either direction. The measure of DFC variation has been often used to explore the dynamics of specific connections ^29, 30, 31^, and is helpful for understanding which connections contributed the most to altered stability. DFC variation for each ROI was calculated as the standard deviation of DFC across sliding-time windows, and compared between the two states. As shown in Fig. 4B, DFC variation for the left CalS with its neighboring and contralateral regions was larger in movie watching than in resting state. Fig. 4C revealed that DFC variation for the left pMTG with the left OTC and left pFG was smaller in movie watching than in resting state.

### Exploration and validation of the stability measurement

The functional stability reported above was measured in a voxel-to-voxel approach. This approach regarded voxel-level DFC (tens of thousands of voxels) as features to determine the functional architecture of a given voxel, thus incurring a large computational load. We also explored an approach that could reduce the computational load, and determined to what extent the findings obtained with a voxel-to-voxel approach were preserved. We used a voxel-to-atlas approach, in which the features were defined in terms of DFC with 200 parcellations from the atlas created by Craddock and colleagues ^32^, and stability of a given voxel was computed as KCC of DFC between that voxel with all parcellations (see Supplementary Note 1). For the first dataset, the profile of intrinsic stability derived using the voxel-to-atlas approach was very similar to that derived using the voxel-to-voxel approach (Fig. S3). Statistically, across-subject correlation analyses (n = 216) revealed extremely high correlation between voxel-to-atlas KCC and voxel-to-voxel KCC for all measured voxels (mean r = 0.921, range from 0.728 to 0.975), indicating that the voxel-to-atlas KCC explained most variance of the voxel-to-voxel KCC. For the second dataset, the voxel-to-voxel and the voxel-to-atlas approaches also produced similar results of stability modification by movie watching (Fig. S4).

We next explored whether our main results were specific to the sliding-window parameters. DFC and then KCC were recomputed, with other settings of window length, window step, and window type (see Supplementary Note 2). One-sample T-tests of intrinsic stability revealed that results were consistent across different settings (Fig. S5), indicating that our main results were not impacted by the sliding-window parameters. In addition, contrasts of stability between movie watching and resting state using several different sliding-window parameter settings revealed similar patterns of modification in functional stability, especially for the OTC and extended occipital areas (Fig. S6).

## Discussion

The brain’s functional organization changes dynamically even during rest ^21, 33^. While prior studies have explored temporal variability or flexibility of functional organization ^7, 8, 34^, this study investigated the other side, the stability of functional architecture that may represent a critical property of the brain ^1^. We first characterized how stability of functional architecture is distributed across the brain, and found the apex of functional stability over time in high-order association regions, especially those in the DMN, rather than unimodal regions. Then we explored how functional stability was modified during natural viewing, and showed that compared to resting state, functional stability during movie watching was increased in high-order visual regions (bilateral OTC, right STS, and left IPS), and decreased in low-order visual regions (bilateral posterior and medial occipital lobes) and the mPFC.

### Stability of functional architecture differs across brain regions

The study on brain dynamics by Allen et al. (2014) clustered highly-structured reoccurring connectivity patterns into sub-states ^35^, suggestive of stability for the brain’s functional architecture, in addition to flexibility. Our results revealed a high level of stability in high-order association regions, especially in the DMN regions (mPFC, AG, and PCC) which had the most stable functional architecture. The PCC and mPFC are considered as the core DMN regions involved in internally-directed thought ^36^, and DMN connectivity has been associated with consciousness ^37^. As a part of the DMN, the AG is also proposed to subserve convergence of multisensory information, and to thus participate in various complex tasks ^38^. The DMN regions situate at one end of a principal gradient of brain functional organization, of which the other end is anchored by primary sensory and motor regions ^39^. Previous studies observed medium or low flexibility of functional architecture in core DMN regions ^7, 8^. Importantly, these two studies did not characterize the core DMN regions as high-order association regions. These regions do not specifically process signals of one modality, and are generally considered to be brain hubs conducting high-order cognitive processes ^5, 6, 36^. The high stability may provide a foundation for the DMN regions to integrate multimodal information over a long time scale.

Other regions with high functional stability were mainly located in the FPN and VAN, including the DLPFC, AIns, and SMG. The DLPFC plays a critical role in executive functions which refer to high-order organization and dynamic tuning of behaviors and thoughts ^40, 41^. The AIns has been linked with human awareness ^42^, and with an integral hub for high-order cognitive control ^43^. On average, functional architecture appeared less stable in the FPN and VAN than in the DMN, which was probably due to greater DMN activity during resting state ^19^. The present study extends previous findings by showing that the feature of global connection for these high-order association regions ^5, 6^ is stable over time within a state. This observation is contrary to the hypothesis that association regions change their functional connections frequently since they switch to interact with distributed regions of different modalities.

In comparisons across the brain, functional stability in sensory-motor cortices was much lower than that in high-order association regions, indicating that unimodal regions reorganized their activity or connection patterns over time. Unimodal regions accumulate information in a short time scale ^11^, so their functional organization is not necessary to be stable over time. Moreover, neural activity of unimodal regions is driven by both external stimuli and top-down modulation from high-order regions ^44^, and external dependence may explain the decreased functional stability (see below).

### Stability of functional architecture differs between states

During movie watching, viewers receive a sequence of visual images which constantly change in form but exhibit coherence in meaning. This task thus gives rise to a continuous and natural state, as compared to conventional experimental tasks with discrete and independent events and stimuli. Our results revealed that compared to resting state, movie watching decreased functional stability of the bilateral primary visual cortices and mPFC, and increased functional stability of bilateral OTC, left IPS, and right STS which support high-order visual processing. The primary visual cortices are proposed to process the form of visual images (e.g., orientation, color, etc.) ^45, 46^. Since sensory inputs directly affect neural activity of these regions, the decreased stability could possibly be explained by adjustment of functional architecture to the changes of received visual form over time. Considering that the primary visual cortices also receive top-down influence from the high-order regions of visual stream and top-down control from the frontoparietal regions ^47, 48^, another possibility is that the switch of connections to these regions caused the reduction of functional stability. The analysis of ROI-based DFC variation comparison lent support to the former explanation, which revealed larger DFC variation to neighboring regions within the visual cortices.

In contrast, increased functional stability during movie watching was found in regions that participate in high-order visual processing ^46, 49^. In the ventral visual stream, the posterior MTG contributes to visual motion processing ^50^, while the STS and OTC are considered to integrate auditory and visual information ^17, 18^. In the dorsal visual stream, the IPS participates in visual processing due to its role in attention and space processing ^51, 52^. To derive a comprehensive perception and cognition of sight, high-order visual regions not only process visual information alone, but also integrate information from other modalities ^53, 54^. Movie watching requires accumulation of audiovisual information and integration of multimodal information over time. Accordingly, as shown by our results, the functional architecture for these regions did not change to a large extent over the course of movie watching, but was fairly stable. Interestingly, although both the primary and high-order visual cortices were recruited by movie watching (Fig. 5) ^27^, they could be distinguished by the direction in which functional stability was modified by the task, suggestive of significance for this measurement. On the contrary, the functional stability of the mPFC, a high-order region, appeared to reduce during movie watching. This region integrates information over a rather long window ^11^. A 10-min movie clip may be not long or integral enough to elicit a stable connectivity pattern for the mPFC. Future studies using complete versions of movies can address this issue.

### Significance of the functional stability measurement

The distribution and modification pattern of functional stability illustrates from a dynamic view how functional organization adapts to fulfill a complex naturalistic task. The functional architecture of unimodal regions changed with alterations of explicit forms of the input, while stability of the functional architecture of high-order regions allows neural integration both across modalities and across time. This distinction is in line with the previous finding that a hierarchy of temporal scales to integrate information exists in the visual system ^3^. It also echoes the resting state finding that functional architecture appears more stable in high-order association regions than in unimodal regions, since functional architecture in resting state is considered as a composite reflection of multiple task states ^4^.

We speculate that high functional stability in association regions may render the brain adaptive to the environment. During conscious processing, the brain selects information for global broadcasting ^1^, which should be carried out by high-order association regions through their distributed functional connections ^5^. Our findings thus provide evidence of a neurobiological basis from the functional network perspective for the stability property of the brain. A variety of complex cognitive functions require the brain to coordinate information from multiple modalities over time ^2, 4^. So far, it has remained largely unclear whether association regions organize functional architecture in a stable or a flexible manner, to perform integration processes within a continuous state. The present study provides strong evidence for stability. High stability within a state as we found does not contradict high flexibility between tasks or states observed in prior studies ^13, 14^. The stability property (without frequent alteration of connectivity) may provide the efficient capacity to coordinate information over time.

### Methodological consideration and implication for future studies

Here we measured functional stability using a voxel-to-voxel approach. Differences in data analytic approaches may explain inconsistencies between our findings and previous ones. The studies by Zhang et al. (2016) and Yin et al. (2016) found high flexibility for high-order association regions ^7, 8^, while we found high stability for these regions. Those studies employed the AAL atlas and analyzed data in an atlas-to-atlas approach. The AAL atlas separates the brain into 90 functionally inaccurate parcellations that cannot adequately reflect the functional architecture of the brain ^12^. Such analyses would result in an imprecise estimation (Fig. S7). For future studies, we therefore recommend using a refined division of the brain (e.g., voxel-level) to define functional architecture of the brain and examine derived measurements. The voxel-to-atlas approach yielded a pattern of results similar to that using the voxel-to-voxel approach, so when computational resources are limited, the voxel-to-atlas approach is also admissible.

Several issues can be further addressed in the future. First, as a critical feature of the brain, the stability of intrinsic functional architecture and the extent of its modification by naturalistic tasks can be taken as potential biomarkers for quantitative diagnoses of mental disorders. For example, patients with major depression disorder could show less modification of stability when engaging in a task, which is associated with mental slowing. Second, we were unable to accurately quantify functional stability of regions near cavities. Future studies using scanning sequences that increase signal-to-noise ratio for regions near cavities will be required to address this issue.

In conclusion, the functional architecture of high-order association regions is stable over time within a continuous state, and functional stability of this type of regions is increased when they are employed in a task, suggestive of their role in coordinating neural information from successive moments. By contrast, unimodal regions vibrate their functional architecture to process ever-changing stimulus forms. The division of labor between these two types of regions may reflect the way in which the human brain implements high-level cognitions.

## Methods

### Data sources and participants

Two open neuroimaging datasets were used in the present study. The first was obtained from the CoRR (Consortium for Reliability and Reproducibility) release ^20^. To keep scanning parameters (e.g., TR) and instructions uniform across subjects, only one site with the largest sample size was used, which contained resting-state fMRI data of 216 young adults (104 females; mean age = 20.0 years, range: 17 – 27 years). The resting-state scanning lasted for 8 min 2 s during which participants were asked to remain still and think of nothing specifically, with their eyes open. For the second dataset obtained from the HBN (Healthy Brain Network) released by the Child Mind Institute, fMRI data were acquired for 32 children and adolescents (20 females; mean age = 12.1 years, range: 7 – 19 years) while they were at rest and while they watched an audiovisual movie ^25^. There were two runs of resting-state scans each lasting 5 min, and a run of movie watching. The movie was a 10-min clip of an animated film named “Despicable Me” (exact time from 1:02:09 – 1:12:09).

### Data preprocessing

We used Matlab-based toolboxes of SPM12 and DPABI to run data preprocessing ^55^. For the first dataset, the initial 10 functional volumes (20 s) were deleted to allow for signal stabilization. Functional images were corrected for slice acquisition timing differences and head motion. Nuisance covariates, including linear trend, Friston 24 head motion parameters, white matter signal, and cerebrospinal fluid signal, were regressed out from the functional signal. Then the functional images were normalized to MNI space by DARTEL. Band-pass temporal filter (0.01 – 0.1 HZ) and spatial smoothing (6 mm FWHM kernel) were applied to the normalized functional images. For the second dataset, we also preprocessed the functional imaging data following the above procedure except that the initial 25 volumes (20 s) were removed. In addition, slice timing correction was not conducted, since this dataset employed a multiband scanning series and the repetition time (0.8 s) was short. We used the same procedure to preprocess data of the movie-watching run and the resting-state runs, to make them comparable. Subjects with maximum head motion larger than 3 mm in displacement or 3° in rotation were excluded from subsequent analyses, as well as those with mean frame-wise displacement (FD) larger than 0.25 mm. Overall, 16 subjects for the first dataset and 83 subjects for the second dataset (children and adolescents generally have larger head motion during scanning) were excluded. For the remaining 32 subjects of the second dataset, head motion (mean FD) did not differ significantly between the movie-watching run and the resting-state runs (p = 0.241).

### Computation of stability of dynamic functional architecture

For a voxel in the brain, the stability of functional architecture was defined as the concordance of DFC over time of that voxel with the whole brain. DFC was calculated using a sliding-window approach, with the window length being 64 s (32 TRs for the first dataset and 80 TRs for the second) and the sliding step being 4 s ^56^. We conducted analyses in a voxel-by-voxel approach, such that DFC was computed between a voxel with all other voxels within the mask, resulting in DFC maps across the 101 time windows for that voxel (Fig. 1). The Kendall’s coefficient of concordance of these DFC maps with time windows as raters was computed as:

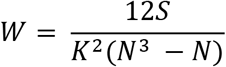

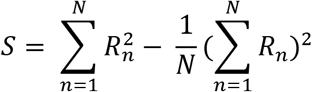

where K is the number of windows, N is the number of connections of that voxel with all voxels within the mask, and R_n_ is the sum of rank for the n-th connection across all windows. For each window, connections are ranked across all voxels based on their functional connectivity strength. W (ranges from 0 to 1) quantified stability of functional architecture of that voxel. The connections of that voxel to the whole brain are regarded as features to represent its functional architecture. Analyses were confined to a grey matter mask, which was created by thresholding the mean grey matter density across participants at 0.2 and intersected with a group mask of 90% coverage of functional images. The derived KCC was z-standardized across the grey matter mask, to increase comparability across participants and conditions.

### Characterization of intrinsic functional stability across the brain

For the first dataset, one-sample T-tests on the KCC z-score were conducted across the group mask, with age, sex, and head motion (mean FD) as covariates. In addition to showing the profile of stability across the brain, we also computed the ratio of voxels with positive and negative KCC after multiple comparison correction using Gaussian Random Field (GRF) theory (with voxel p < 0.001 and cluster p < 0.01, two-tailed; the same below), for each of the seven brain networks ^24^.

To examine whether functional stability was also higher in high-order association regions than unimodal regions within the visual network in the left hemisphere, we selected four unimodal regions of interest (ROIs) located in the primary visual cortex and six high-order association ROIs including V3, V4, and four MT regions (Fig. 3). Their coordinates were the same as those used in the study by Yeo et al. ^24^, and for each ROI, a sphere centered on the coordinates was created with a radius of 4 mm. Functional stability was averaged across the ROIs, for the high-order visual regions and the unimodal visual regions, respectively. Paired-sample T-tests were conducted to compare the averaged functional stability between these two types of visual regions. We also examined the regions in the right hemisphere that were contralateral to the above ROIs.

In addition, to examine whether functional stability was greater than expected by random, observed KCC was compared to one derived from simulated data. The preprocessed functional images of a whole run were transformed to the frequency domain using FFT, and for each voxel the phases of frequency bands were randomized, with the amplitude unchanged. This method removed the temporal alignment of neural signals but kept the amplitude, and thus resulted in a stochastic baseline for the measurement. The KCC was compared between observed data and simulated data for each voxel with paired-sample T-test, using raw values instead of z-scores.

### Modification of functional stability during task state

For the second dataset, since the duration of the movie run was twice the duration of the two resting-state runs, we divided the movie run into two parts and deleted the beginning 20 s from the latter part of the movie run. This resulted in a duration of 280s for each part of the movie run, equal to that of the resting-state runs. For each participant, voxel-to-voxel KCC was computed, z-standardized, and then averaged for the two resting-state runs and for the two parts of the movie run, respectively. The averaged KCC z-score was compared between movie watching and resting state with paired-sample T-tests. GRF theory was used to correct for multiple comparisons. We used a strict correction criterion (cluster p < 0.01, two-tailed) to avoid inflating false positive rates ^26, 57^.

To derive ISC for a given subject, we correlated the neural activity of that subject to the averaged neural activity of the remaining subjects in each voxel. Then the Fisher’s transformation was applied to the correlation coefficient. ISC was computed for all subjects in this way. At the group-level analysis, the ISC was compared to zero using one-sample T-test across the brain, and the mean ISC was also computed. GRF theory was applied to corrected for multiple comparisons. Based on previous research, we also used a threshold of r > 0.25 to eliminate regions with a low level of ISC ^27^.

## Acknowledgments

The authors appreciate the editorial assistance and support of Dr. F. Xavier Castellanos. This work was supported by the National Key R&D Program of China (2017YFC1309902), the National Natural Science Foundation of China (81671774 and 81630031), the Hundred Talents Program of the Chinese Academy of Sciences, and Beijing Municipal Science & Technology Commission (Z161100000216152).

## Author contributions

L.L. and B.L. acquired and preprocessed the data. L.L and C.Y. designed the research and analyzed the data. All authors interpreted the results and wrote the manuscript.

