## Supplementary Materials for "Stability of dynamic functional architecture differs between brain networks and states"

### 1. Comparison of the voxel-to-voxel and the voxel-to-atlas approaches

For the term of “voxel-to-voxel”, it defines the precision levels for the unit of measurement and for the features used to derive it. The former “voxel” refers to the resolution of measurement in the brain, in which functional stability is computed one by one voxel. The latter “voxel” refers to the resolution of features used to compute the measurement, in that the DFC for a given region is computed with all other voxel across sliding windows and then used to quantify the Kendall’s coefficient of concordance (KCC) for that region. The voxel-to-voxel approach utilized complete information to derive functional stability, and provided a refined characterization of the brain, but was computationally intensive.

To reduce computation load, we speculated that decrease of feature resolution would not lose critical information significantly if the parcellations of the brain using an atlas could capture main variation of signals. Here we used the Craddock’s atlas, in which the brain was divided into 200 subregions via spatially constrained clustering of functional signals [1]. The stability of functional architecture was computed in a voxel-to-atlas approach that decreased the number of features from tens of thousands to hundreds. The voxel-to-atlas DFC computation resulted in a set of DFC vectors (with each element for a subregion in the atlas) across sliding windows for each voxel. KCC of DFC vectors of that voxel was computed with window as raters, and converted to z-scores. The results generated using this approach were present in Fig. S3 for the profile of intrinsic functional stability and in Fig. S4 for the modification of functional stability by movie watching, revealing similar patterns as those reported using a voxel-to-voxel approach.

For the first dataset, we computed across-subject correlation for each voxel and across-brain correlation for each subject between voxel-to-atlas KCC and voxel-to-voxel KCC. The across-subject correlation was extremely high across the brain (mean $r = 0.922$, range from 0.732 to 0.975 for the first session; and mean $r = 0.920$, from 0.728 to 0.972 for the second session). The across-brain correlation was also high for all participants (mean $r = 0.941$, from 0.776 to 0.986 for the first session; mean $r = 0.938$, from 0.805 to 0.985 for the second session). This result indicated that the voxel-to-atlas KCC explained most variance of the voxel-to-voxel KCC. As the voxel-to-atlas approach cost much less computational resources (only about one hundredth computing time), it was more suitable for the cases where computational resources were limited or for exploratory aims.

**2. Verification of results with different sliding-window parameter settings**

The sliding-window approach has been used and developed for neuroimaging studies only in recent years, and therefore it remains controversy about its parameter setting [2]. Window length defines the duration of consecutive substates, and is related to the number of data points in a window used to compute functional connectivity (FC). The duration should be at least 30-60 s long to get a robust estimation of FC for a cognitive state [3]. We thus used 64 s (32 data point for the first dataset and 80 data point for the second dataset) as the window length in the analyses reported in main text. Here we further rerun the analyses with three different window length, that was 32 s, 48 s, and 128 s, respectively, to examine whether our results were influenced by specific window length. As shown in Fig. S5A, the pattern of functional stability during resting state was highly consistent across different window length.

Window sliding step defines the shift time between two successive windows, and is also negatively related to the number of raters for the KCC computation. We rerun the analyses with a step of 2 s (instead of 4 s used in the main text). The results showed a similar pattern of functional stability across the brain as those using other sliding-window parameter settings (Fig. S5B). Window type defines the shape of window and the function to extract BOLD time series for each windowed segment from data. We then rerun the analyses using the Hamming window (instead of rectangular window used in the main text), and the pattern of functional stability almost remained unchanged (Fig. S5C). Taken together, the consistent pattern of functional stability of the brain across different sliding-window parameters indicated the robustness of the results.

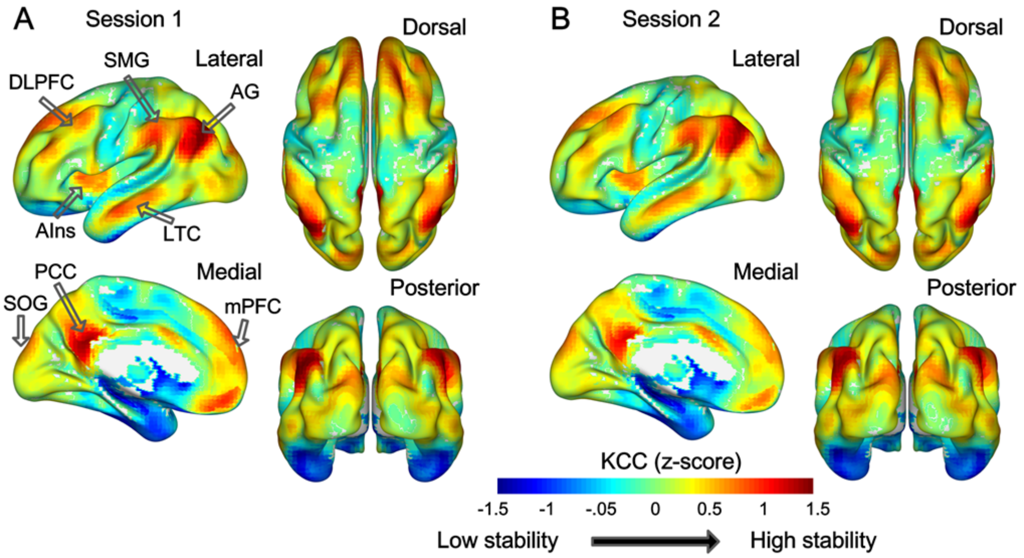


Figure S1. Averaged functional stability (converted to z-scores, mean values instead of one-sample T values in Figure 2) in resting state for session 1 (A) and session 2 (B). The pattern of high vs. low stability across the brain is similar to the pattern of one-sample T-tests in the main text. High stability was observed in several association regions indicated by black hollow arrows. KCC, Kendall's concordance coefficient; DLPFC, dorsolateral prefrontal cortex; AG, angular gyrus; AIns, anterior insular; LTG, lateral temporal cortex; SOG, superior occipital gyrus; PCC, posterior cingulate cortex; mPFC, medial prefrontal cortex.

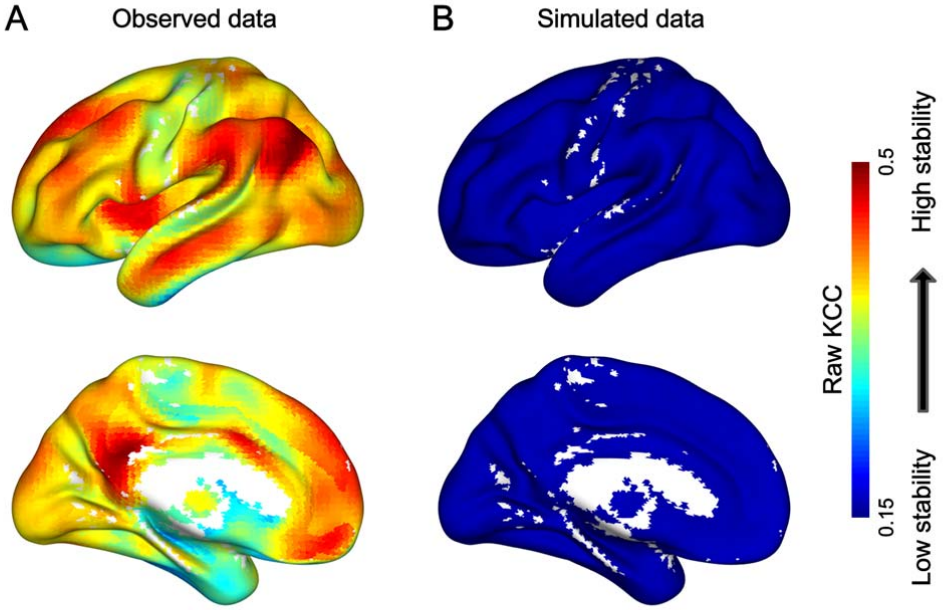


Figure S2. Functional stability in the form of raw value derived from real data (A) and simulated data (B). A large value (red) denotes high levels of stability. To derive the simulated data, the real data were transformed to the frequency domain using FFT, and for each voxel the phases of frequency bands were randomized, with the amplitude unchanged. Stability derived from simulated data severed as a null distribution for testing whether functional stability of a region is higher than random level.

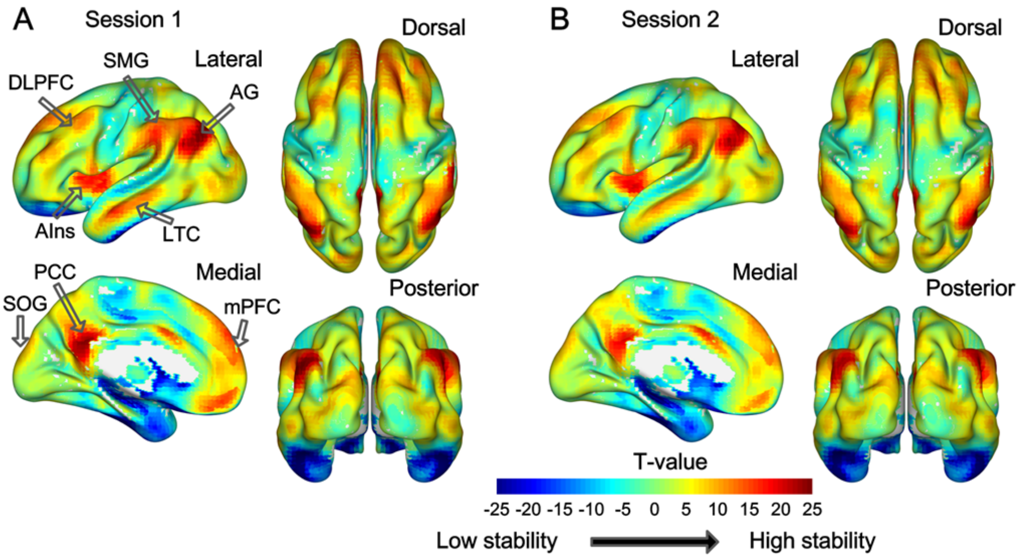


Figure S3. Results of one-sample T-tests on functional stability in resting state using the voxel-to-atlas approach. The pattern of high vs. low stability across the brain is similar to that derived using the voxel-to-voxel approach reported in the main text (Fig. 2), for both session 1 (A) and session 2 (B). For the abbreviations, please refer to the legends of Fig. S1.

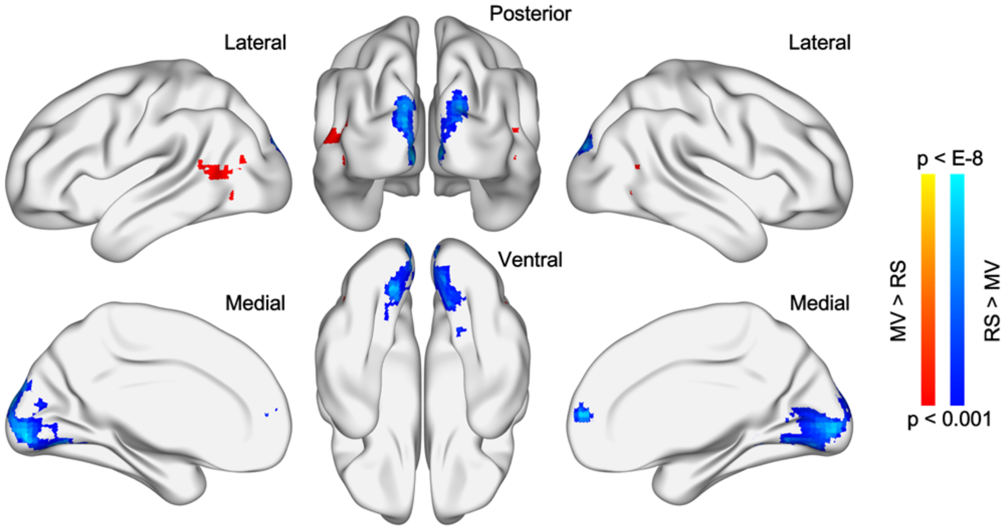


Figure S4. Results of paired-sample T-tests on functional stability between movie watching and resting state using the voxel-to-atlas approach. The pattern of state difference in stability is similar to that reported in the main text (Fig. 3). MV, movie watching; RS, resting state.

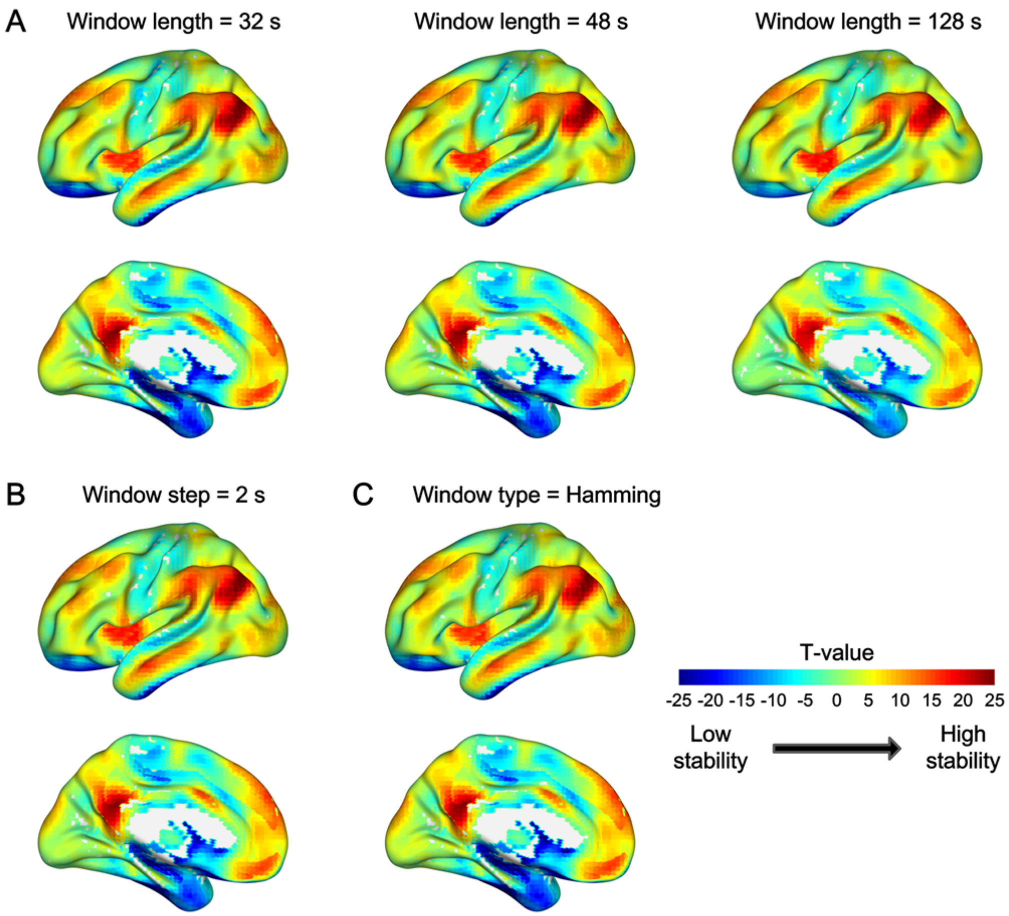


Figure S5. Results of one-sample T-tests on functional stability in resting state using different sliding-window parameters. The window length was set to 32 s, 48 s, and 128 s, respectively (A), other than 64 s. The window sliding step was set to 2 s (B), other than 4 s. The Hamming window was used (C) instead of the rectangular window. The results were consistent across different sliding-window parameter settings.

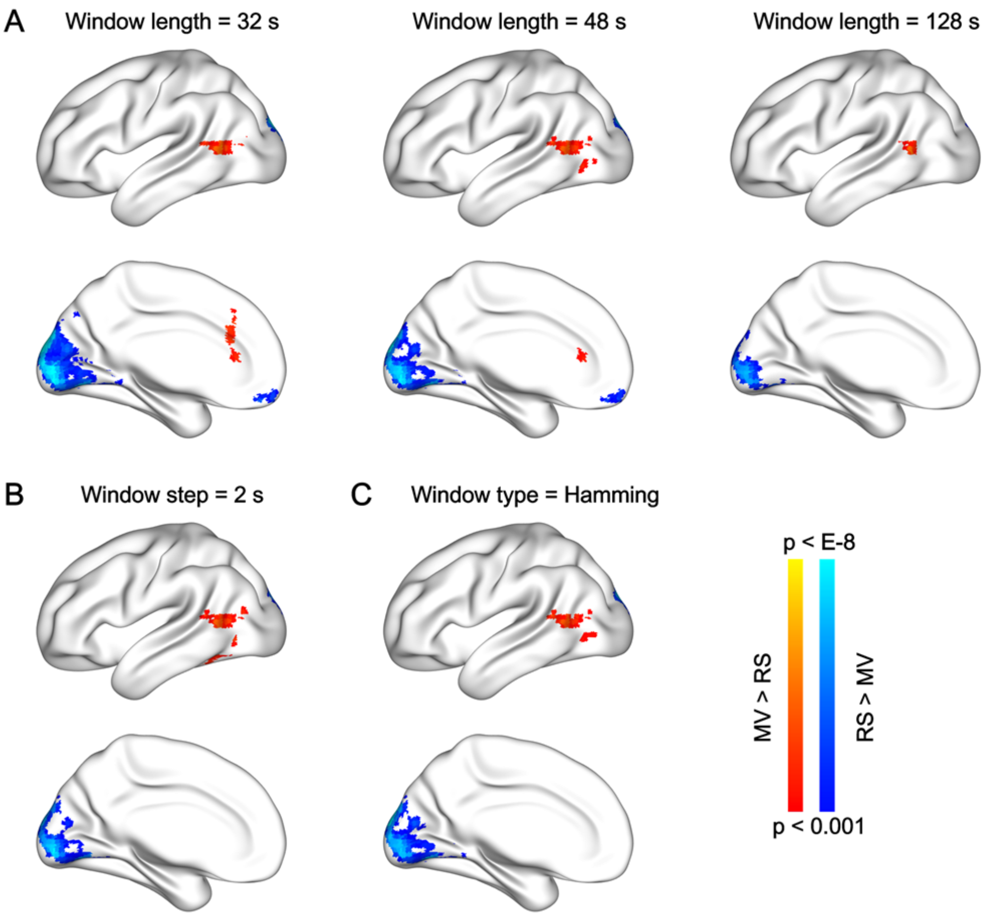


Figure S6. Results of paired T-tests on functional stability between movie watching and resting state using different sliding-window parameters. The window length was set to 32 s, 48 s, and 128 s, respectively (A), other than 64 s. The window step was set to 2.4 s (B), other than 4 s. The Hamming window was used (C) instead of the rectangular window. The results were similar across different sliding-window parameter settings.

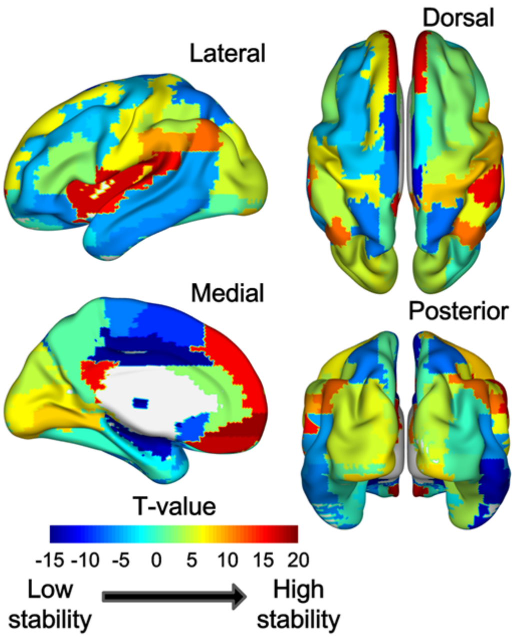


Figure S7. Results of one-sample T-tests on functional stability in resting state computed using the atlas-to-atlas approach. The atlas employed was the Automated Anatomical Labeling (AAL) atlas. This analysis was conducted in order to reveal how such a approach would produce an inaccurate estimate of functional stability.
